# Plasma membrane damage removal by F-actin-mediated shedding from repurposed filopodia

**DOI:** 10.1101/2019.12.23.887638

**Authors:** Shrawan Kumar Mageswaran, Wei Y. Yang, Yogaditya Chakrabarty, Catherine M. Oikonomou, Grant J. Jensen

## Abstract

Repairing plasma membrane damage is vital to eukaryotic cell survival. Membrane shedding is thought to be key to this repair process, but a detailed view of how the process occurs is still missing. Here we used electron cryotomography to image the ultrastructural details of plasma membrane wound healing. We found that filopodia-like protrusions are built at damage sites, accompanied by retraction of neighboring filopodia, and that these repurposed protrusions act as scaffolds for membrane shedding. This suggests a new role for filopodia as reservoirs of membrane and actin for plasma membrane damage repair. Damage-induced shedding was dependent on F-actin dynamics and Myo1a, as well as Vps4B, an important component of the ESCRT machinery. Thus we find that damage shedding is more complex than current models of simple vesiculation from flat membrane domains. Rather, we observe structural similarities between damage-mediated shedding and constitutive shedding from enterocytes that argue for conservation of a general membrane shedding mechanism.

## Main

Maintaining the integrity of the plasma membrane is critical for the survival of eukaryotic cells, and cells exhibit a rapid response to plasma membrane injury. Injury may include mechanical damage, chemical insults or the introduction of foreign pore-forming proteins such as the bacterial toxin streptolysin O (SLO) or the perforin secreted by cytotoxic T lymphocytes and natural killer cells^1–3^. Laser ablation of the plasma membrane elicits similar responses^2^. Following damage, a rapid influx of Ca^2+^ and ensuing exocytosis of lysosomes occurs^4–6^ and the membrane is resealed within a few seconds to a few minutes. Several models, not mutually exclusive, have been proposed for how resealing occurs: (1) in what is known as the “patch model”, a damaged region is replaced by fusion of an intracellular vesicle with the plasma membrane, sloughing off the damaged membrane in the process^7^; (2) in the endocytic model, damaged regions are internalized by endocytosis^3, 8^; and (3) in the shedding model, damaged regions bud from the plasma membrane and are shed as extracellular vesicles^2^. While the patch model has not yet been validated experimentally, evidence exists for both endocytosis and shedding. For example, SLO-induced pores have been shown to be eliminated by both endocytosis^9^ and shedding^1, 10, 11^. Several pieces of recent evidence suggest that shedding may be the dominant mechanism for plasma membrane resealing. First, repair occurs even at low temperature, a condition that stalls other mechanisms, including endocytosis^12^. Second, after SLO treatment, SLO-containing vesicles are released in an annexin-dependent process^10, 11^. Third, extracellular vesicle-like structures are observed by scanning electron microscopy (SEM) at sites of laser damage^2^. And finally, <u>E</u>ndosomal <u>S</u>orting <u>C</u>omplexes <u>R</u>equired for <u>T</u>ransport (ESCRT), known to be involved in membrane remodeling processes similar to shedding, were found to be important for resealing small holes (<100 nm)^2^.

Shedding is a ubiquitous process with diverse functions apart from plasma membrane damage repair. Depending on the cell type, shedding may be constitutive, and is amplified by environmental stimuli^13^. Shed vesicles can deliver enzymes to the intestinal lumen for digestion^14^ and detoxification of bacterial lipopolysaccharides^15^ to control bacterial population^16^. Vesicles released by neutrophils and other immune-response cells regulate inflammation^17–19^. Vesicles from astrocytes and neurons contain signaling growth factors^20, 21^. Vesicles of vascular origin play important roles in angiogenesis, metastasis, atherothrombosis and other diseases^22–24^. Despite its importance, the molecular mechanism(s) of shedding remains unclear. The process is best understood in the microvilli of gut enterocytes where central actin bundles are connected to the plasma membrane by spoke-like radial densities of the motor protein Myo1a^25^. Myo1a is important for stabilizing the microvillus, as well as promoting shedding at the distal tip (possibly by propelling membrane over actin bundles^26^).

Here we investigated the ultrastructural details of plasma membrane repair in HeLa cells following laser damage. By electron cryotomography (cryoET), we observed that cells relocate actin and membrane to sites of damage to generate F-actin-rich filopodia-like protrusions that act as scaffolds for vesicle shedding. When N-WASp (required for actin nucleation) or Myo1a was disrupted, cells displayed defects in generating protrusions and shedding vesicles, and disruption of Vps4B, an ESCRT AAA ATPase previously shown to be important for wound repair^2^, led to defects in membrane scission, the final step in shedding. These results reveal that shedding in response to plasma membrane damage is strikingly similar to constitutive shedding from microvilli, suggesting a common underlying mechanism.

## Results

### Membrane and actin are relocated to sites of damage

We grew HeLa cells expressing CHMP4B-EGFP, an ESCRT-III protein that gets recruited from the cytosol to the sites of plasma membrane injury, on glass and induced damage with a 640 nm laser. Compared to other damage methods including detergents, toxins and mechanical disruption, laser treatment offers reproducibility and precise control over the position of damage sites. It also offers the convenience of administering the insult and monitoring the response with the same instrument, in our case a scanning confocal microscope. We adapted a previously published scheme for laser-based (UV wavelength) damage^2^ by introducing a photosensitizer that causes damage by generating reactive oxygen species (ROS) through fluorescence emission^27–29^. Photosensitizers can be used to specifically damage target cellular compartments such as mitochondria^30^ and lysosomes^31^. For our workflow, we preincubated cells with the plasma membrane-localized photosensitizer disulfonated aluminum phthalocyanine (AlPcS2a).

We first standardized the parameters for laser damage. We damaged a 3 µm-wide circular area lacking (by bright-field imaging) filopodia on the edge of a cell. We observed that prolonged laser illumination caused numerous blebs – bulging regions of plasma membrane formed by reorganization of cortical F-actin – around the cells (Supplementary Movie 1 – cell 1). We therefore calibrated the laser illumination by reducing the number of pulse cycles (see Materials and Methods) such that blebbing was limited and cells recovered while remaining attached to the glass surface (Fig. 1, Supplementary Movie 1 – cells 2-6). 150 illumination cycles of the laser produced more reproducible results compared to 100. In total, we imaged the response to laser damage in ∼50 cells, with the results summarized in Table 1. Consistent with previously published data^2^, we observed that CHMP4B-EGFP went from being diffusely cytosolic to accumulating at damaged sites (in 40/48 cells) at 5-15 minutes post-damage and reproducibly persisted for at least 40 minutes (Fig. 1). This recruitment of CHMP4B-EGFP was specific to damage sites, compared to randomly chosen control regions nearby (Fig. 1 – viii; note that the control regions Sq2 and Sq3 show some decrease in intensity due to relocation of cytosolic protein and/or photobleaching).

**Table 1.** Live-cell microscopy of cells after laser damage.

| Laser Damage | CHMP4B-EGFP recruitment | Blebbing | Retraction of blebs | Loss of filopodia | Appearance of at least 1 new protrusion | Partial retraction of cell area | Actin relocation |
| --- | --- | --- | --- | --- | --- | --- | --- |
| **150 pulse cycles of laser damage** |  |  |  |  |  |  |  |
| Damaged | 22/26 | 21/26 | 8/21 | 22/26 | 20/26 | 3/26 | 10/11 |
| Control | 0/25 | 0/25 | - | 4/25 | 17/25 | 0/25 | 0/2 |
| **100 pulse cycles of laser damage** |  |  |  |  |  |  |  |
| Damaged | 18/22 | 11/22 | 5/11 | 13/22 | 6/22 | 3/22 | 4/7 |
| Control | 0/7 | 0/7 | - | 0/7 | 2/7 | 0/7 | 0/3 |
| **Total** |  |  |  |  |  |  |  |
| Damaged | 40/48 | 32/48 | 13/32 | 35/48 | 26/48 | 6/48 | 14/18 |
| Control | 0/32 | 0/32 | - | 4/32 | 19/32 | 0/32 | 0/5 |

**Fig. 1:**
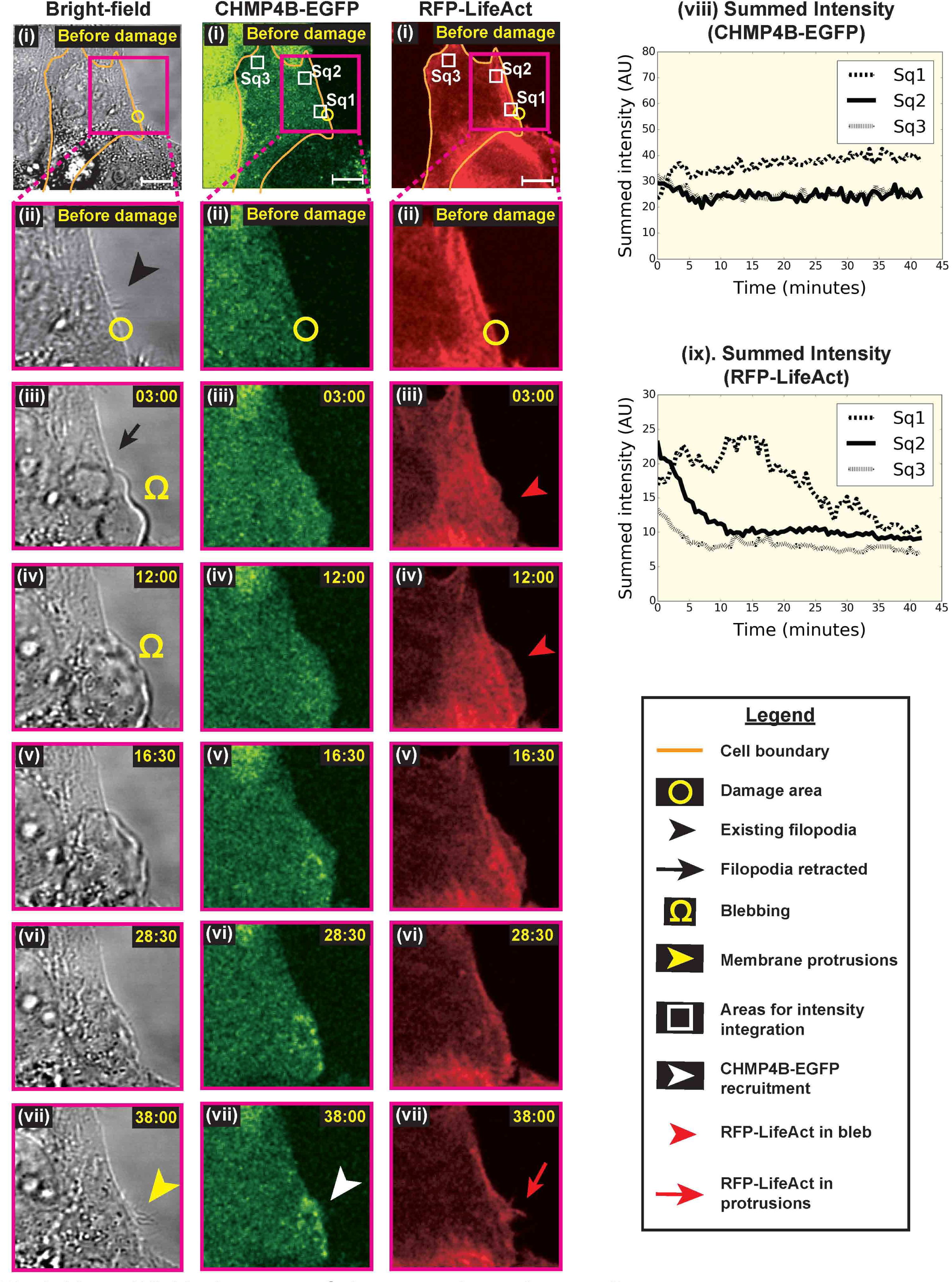
Live-cell light microscopy of plasma membrane damage sites. Light microscopy images of HeLa cells grown on glass (i) before, and (ii-vii) at various time points after damage – (left-to-right) bright-field, CHMP4B-EGFP and RFP-LifeAct imaging. (viii & ix) Integration of pixel intensities for CHMP4B-EGFP and RFP-LifeAct over the square regions marked as Sq1, Sq2 and Sq3 in (i) at various time points after damage. Sq1 lies over the damaged area while Sq2 and Sq3 are control sites. The damage area (yellow circle) is 3 µm in diameter and the time points after damage are denoted in minutes:seconds. Scale bars – 10 µm.

Several bright-field microscopy observations suggested large-scale remodeling of the plasma membrane (Fig. 1a, Supplementary Movie 1 – cells 2-6). First, plasma membrane blebs formed (32/48 cells) and were subsequently retracted (almost completely in 13/32 cells). The extent of blebbing and subsequent retraction varied from one cell to another. Second, dynamic ruffled membrane boundaries appeared (Supplementary Movie 1 – cells 5-6). Third, existing filopodia near damage sites were retracted (35/48). And fourth, new plasma membrane protrusions were formed at the sites of damage (26/48). These protrusions largely resembled filopodia but sometimes displayed pearling, a phenomenon by which tubular membranes spontaneously stabilize into structures with regularly spaced constrictions, resembling beads on a string^32, 33^ (Supplementary Fig. 1). Numerous new protrusions were observed, beginning 10-15 minutes post-damage, and they continued to appear/persist as CHMP4B-EGFP was recruited. The newly formed protrusions often showed punctate CHMP4B-EGFP fluorescence along their lengths (Supplementary Movie 1 – cell 4). The disappearance of existing filopodia and the appearance of new protrusions were sometimes simultaneous and sometimes sequential. Very few control sites in undamaged cells displayed these phenomena (Table 1). Note that we could not quantify the number of protrusions due to the limited resolution of bright-field imaging, so the appearance of (any) new protrusions in control cells likely reflects random rearrangement of a few filopodia. Indeed, the protrusions formed at damage sites qualitatively outnumber the ones at control sites. Finally, a few damaged cells retracted a portion of their area (6/48; Table 1), forming retraction fibers labeled by CHMP4B-EGFP (Supplementary Fig. 2).

Phenomena such as blebbing, plasma membrane ruffling and modulation of cell protrusions are suggestive of a role for F-actin. We therefore imaged damage response in cells labeled with RFP-LifeAct. We indeed observed relocation of F-actin to damage sites (Fig. 1, Supplementary Movie 1 – cells 2-3, and quantified in Fig. 1 – ix). Initially, F-actin was enriched in blebs, consistent with its established role in membrane blebbing^34^. Subsequently, F-actin relocated to the newly formed protrusions.

### CryoET reveals F-actin-rich membrane protrusions and free vesicles at damage sites

To study the ultrastructural details of plasma membrane repair, we used electron cryotomography (cryoET), an imaging technique that can provide high-resolution three-dimensional structural information about biological samples preserved in a near-native frozen-hydrated state. Fortunately, the periphery of adherent HeLa cells is thin enough for direct imaging by cryoET. To target damage sites precisely for cryoET, we used CHMP4B-EGFP recruitment as a marker for cryogenic correlative light and electron microscopy (cryo-CLEM). By light microscopy, we observed that cells grown on EM grids were more susceptible to damage and we therefore reduced the damage area to 1.5 µm in diameter and reduced the number of laser pulse cycles to 35. Also, to standardize the recovery time after damage and prevent cells from undergoing large morphological changes between light microscopy and plunge-freezing, we fixed the cells after damage and recovery, before plunge-freezing. Fixation has been shown to largely preserve ultrastructure^35–37^. As with cells on glass, laser-damaging cells on EM grids resulted in CHMP4B-EGFP recruitment at the damage sites by 10-15 min (Fig. 2a). Based on our earlier results, this time point was also ideal to image new cell protrusions at damage sites. We therefore fixed cells between 10 and 15 minutes post-damage (note that the time window was to accommodate multiple photo-damage experiments on each EM grid). The same damage sites were then precisely located by CLEM before imaging by cryoET. This workflow is summarized in Supplementary Fig. 3, and comprehensively shown through a representative experiment in Supplementary Movie 2. We advise readers to watch this movie before proceeding further.

**Fig. 2:**
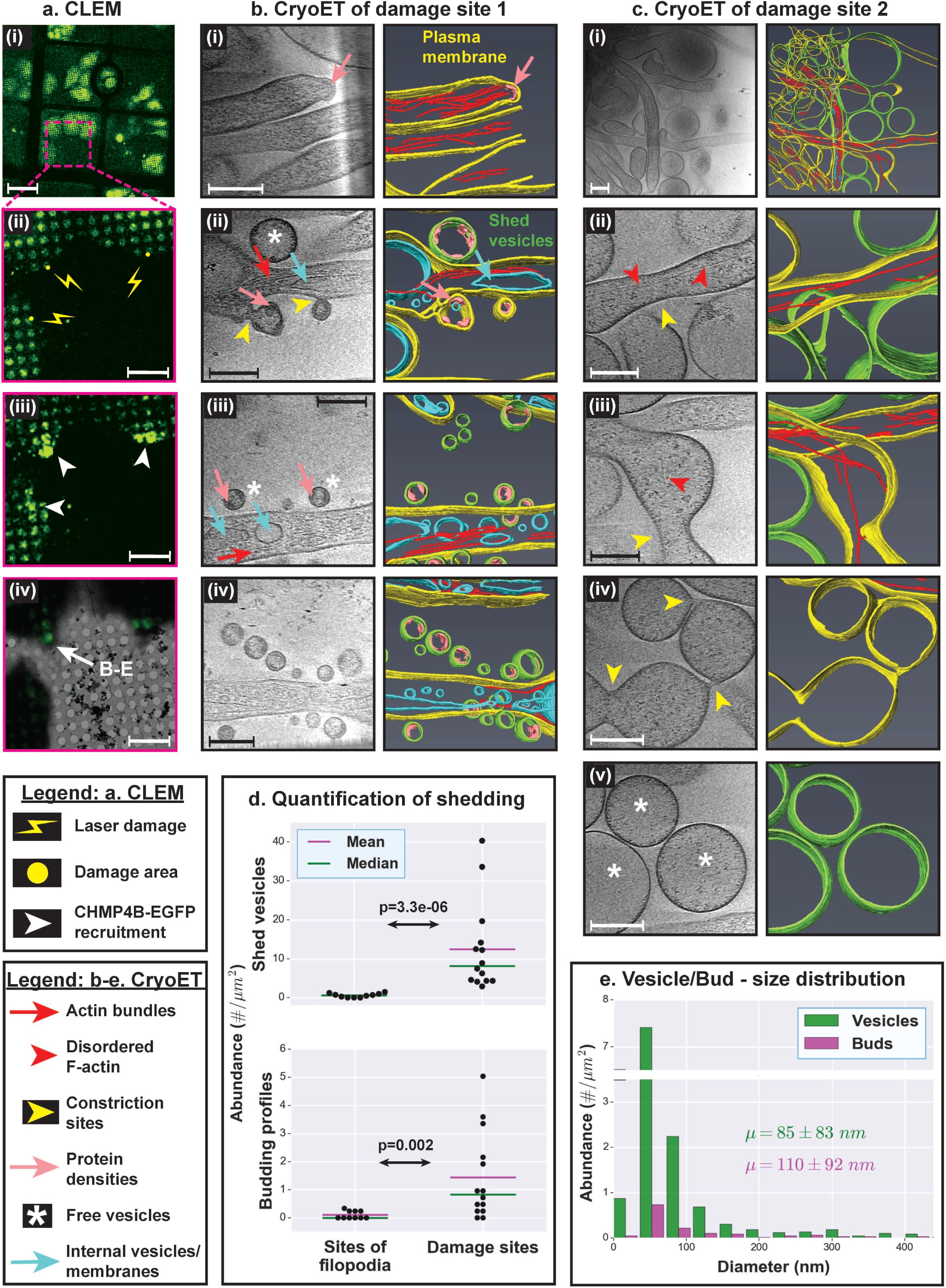
CryoET of plasma membrane damage sites. (a) CLEM. (i) Cells expressing CHMP4B-EGFP grown on an EM grid. (ii) Sites of laser damage, each 1.5 µm in diameter. (iii) Sites of laser damage showing CHMP4B-EGFP recruitment. Cells were monitored for 10 min post-damage and fixed for 45 min using 4% PFA. (iv) Overlay of fluorescence on electron micrograph. (b-c) CryoET of damage sites and corresponding segmentation showing actin-filled plasma membrane protrusions, pearling/budding profiles, shed vesicles and protein densities observed at certain sites of high membrane curvature in budding profiles and shed vesicles. (d) Quantification of shedding. Top panel – abundance of shed vesicles at control sites versus sites of damage (# vesicles per µm^2^ tomogram X-Y cross-sectional area); Bottom panel – abundance of budding profiles at control sites of regular filopodia versus sites of damage (# buds per µm^2^ tomogram X-Y cross-sectional area). Each data point represents a tomogram. In both pairwise comparisons, the distributions for the samples are significantly different from each other as shown by Kolmogorov-Smirnov test (KS test). (e) Size distribution of shed vesicles and budding profiles at damage sites (# vesicles or buds in each size range per µm^2^ tomogram X-Y cross-sectional area) along with the mean ± s.d. values. Scale bars – (a – panel i) 50 µm, (a – panels ii-iv) 10 µm, and (b-c) 200 nm.

CryoET of damage sites revealed numerous F-actin-rich plasma membrane protrusions with budding vesicles, and abundant free vesicles nearby (Fig. 2b-d and Supplementary Movie 2). Budding profiles were even observed on the free vesicles. The size distribution of budding profiles matched that of the free vesicles (Fig. 2e; diameter of 85 ± 83 nm for free vesicles and 110 ± 92 nm for budding profiles; mean ± standard deviation (s.d.)), suggesting that the free vesicles were shed from the protrusions. Further supporting this idea, the budding profiles showed protein densities just inside the plasma membrane similar to those present in the shed vesicles. Free vesicles and budding profiles were found on/near filopodia of control cells as well but were relatively rare (Fig. 2d). A subset of damage site protrusions showed regions devoid of F-actin or with disorganized F-actin, usually at their distal tips (Fig. 2c). Such regions had often lost their tubular morphology and displayed a propensity to pearl, consistent with our light microscopy experiments (note that the pearled regions observed by bright-field microscopy were an order of magnitude larger in size). By comparison, filopodia without damage showed no pearling. Pearled membrane protrusions were pleomorphic, with varying numbers of constriction sites. Free vesicles of comparable sizes were abundant in the vicinity (Fig. 2c), suggesting that pearling-mediated membrane constriction contributes to vesicle shedding.

Damage-induced protrusions resembled canonical filopodia, exhibiting a central bundle of longitudinal F-actin filaments sheathed by plasma membrane (Fig 3a,b). Although the F-actin bundles spanned the entire length of both structures, individual actin filaments were shorter and several filaments were seen originating within the protrusions (consistent with previously characterized filopodia in *Dictyostelium*^38^). Protrusions at damage sites were as abundant as, or in several cases more abundant than, filopodia seen around control cells (Fig. 3c; note that regular filopodia were non-uniformly distributed, so abundance was measured in clusters). Filopodia and damage site protrusions displayed several other striking similarities: (1) similar widths (Fig. 3d); (2) presence of internal vesicles (Fig. 3e), suggesting active membrane trafficking; (3) linker-like densities between filaments and between F-actin and the plasma membrane (Fig. 3a,b – panels ii-iii); (4) branch points, with F-actin bundles at the periphery (Fig. 3a,b – panels iii-iv); (5) F-actin filaments derived from the cortical actin network at the base (Fig. 3a,b – panel iv); (6) similar lengths for linear F-actin filaments (Fig. 3f; 122 nm for filopodia and 163 nm for damage site protrusions); and (7) lateral inter-filament spacing of ∼10 nm (10.2 ± 1.3 nm for filopodia and 10.1 ± 1.2 nm for damage site protrusions; mean ± s.d.). By light microscopy, we also observed recruitment of FusionRed fused to Fimbrin/Plastin-1 (Pls1), an actin-bundling protein found in filopodia, into damage site protrusions (Fig. 3g and Supplementary Movie 3). These results, together with the correlation between the appearance of damage site protrusions and the retraction of neighboring existing filopodia, suggest that the protrusions are repurposed filopodia.

**Fig. 3:**
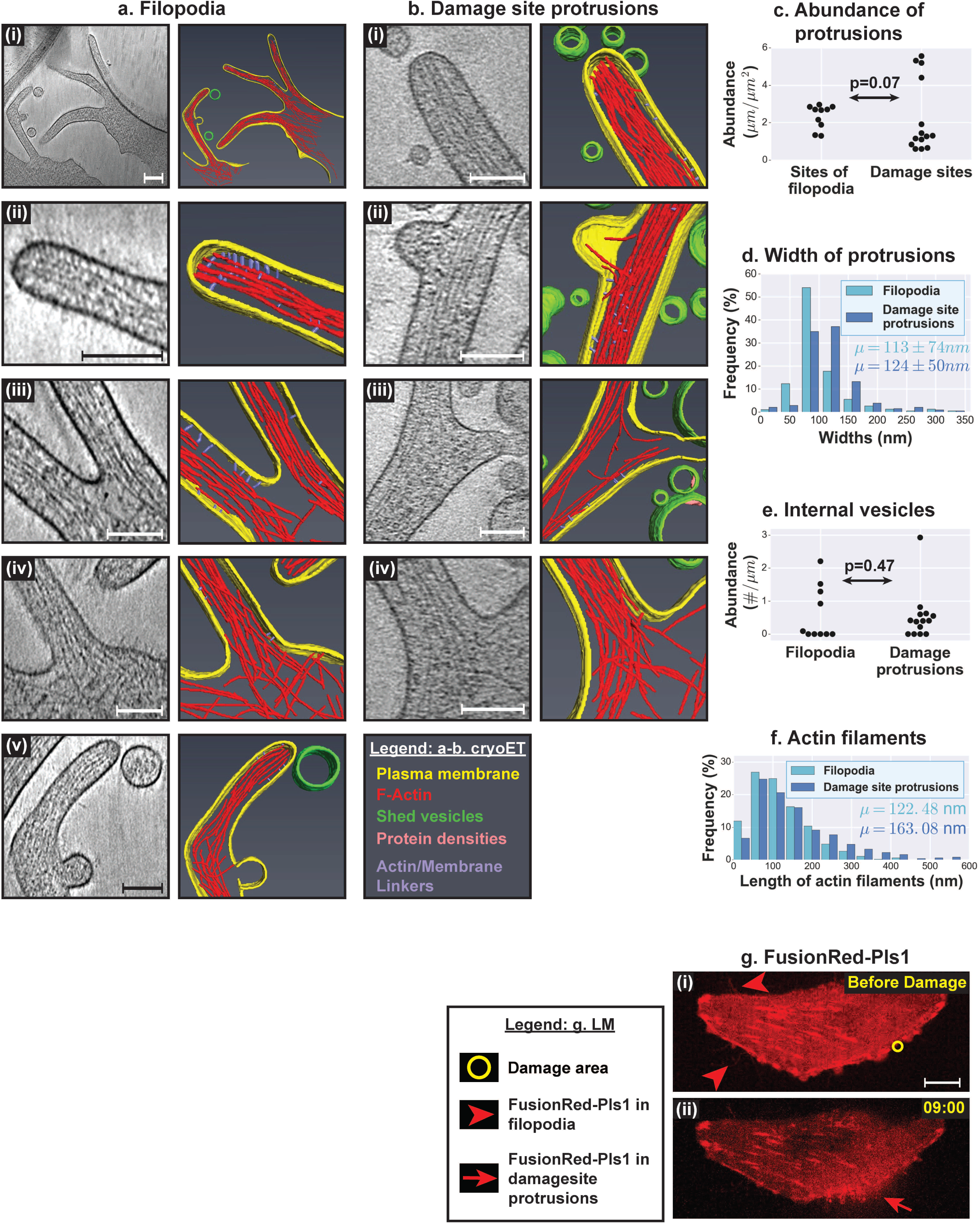
Similarities between filopodia and damage site protrusions. (a-b) CryoET and corresponding segmentation of (a) filopodia at undamaged sites and (b) plasma membrane protrusions at damage sites. (c) Quantification of the abundance of regular filopodia at control sites versus that of plasma membrane protrusions at damage sites (µm total length of protrusions per µm^2^ tomogram X-Y cross-sectional area). (d) Distribution of widths of filopodia versus damage site protrusions (% in each size range) along with their mean ± s.d. values. (e) Quantification of internal vesicles in filopodia versus damage site protrusions (# vesicles per µm of protrusion length). (f) Distribution of lengths of linear actin filaments in filopodia versus damage site protrusions (% in each size range) along with their mean values. (g) Imaging FusionRed-Pls1 in HeLa cells grown on glass (i) before and (ii) after damage. The damage area (yellow circle) is 3 µm in diameter and the time points after damage are denoted in minutes:seconds. In (c) and (e), each data point represents a tomogram and the distributions for the pairs of samples being compared are not significantly different from each other (as shown by KS tests). Scale bars – (a – panel i) 200 nm, (g) 10 µm, and 100 nm for all other panels.

### Actin-rich membrane protrusions are the source of shed vesicles

The abundance of free vesicles and intermediates around damage-induced protrusions strongly suggests that the protrusions act as scaffolds for shedding. In order to test this hypothesis, we analyzed the damage response in cells after disrupting the N-WASp actin-nucleation pathway with wiskostatin. Wiskostatin binds the GTP-binding domain of N-WASp and stabilizes it in an autoinhibited form^39^, thus preventing de novo nucleation of linear chains of F-actin or activation of Arp2/3 to form branched F-actin chains. When cells were treated with wiskostatin for 2-3 hours, before laser treatment, the number of filopodia visible by bright-field imaging was greatly reduced (Fig. 4a – panels I, iii, and v), although the cells still spread on the glass support. We observed an enrichment of vacuole-like vesicles in these cells, consistent with previous observations^40^, but their significance is unknown. Following laser damage, CHMP4B-EGFP was recruited to the damage site as in untreated cells (Fig. 4a – panels ii, iv, and vi). However, no protrusions were visible by bright-field imaging. CryoET similarly showed a significant reduction in the number of damage site-protrusions compared to untreated cells (Fig. 4b,c). Instead, we observed aberrant membrane structures (not found in untreated cells) that may represent accumulation of membranes that failed to form protrusions (Fig. 4b). Shed vesicles were also less abundant than from untreated cells (Fig. 4d,e). Examination of the vesicles revealed that those from wiskostatin-treated cells showed a narrower range of sizes than those from untreated cells, with fewer vesicles larger than 100 nm in diameter (Fig. 4e). This reduction is reflected in the their size distribution (Fig 4d) and their mean sizes (57 ± 44 nm for wiskostatin-treated cells compared to 85 ± 83 nm for untreated cells; mean ± s.d.). In summary, when actin nucleation is blocked, fewer protrusions and vesicles are observed at damage sites, although CHMP4B-EGFP is still recruited.

**Fig. 4:**
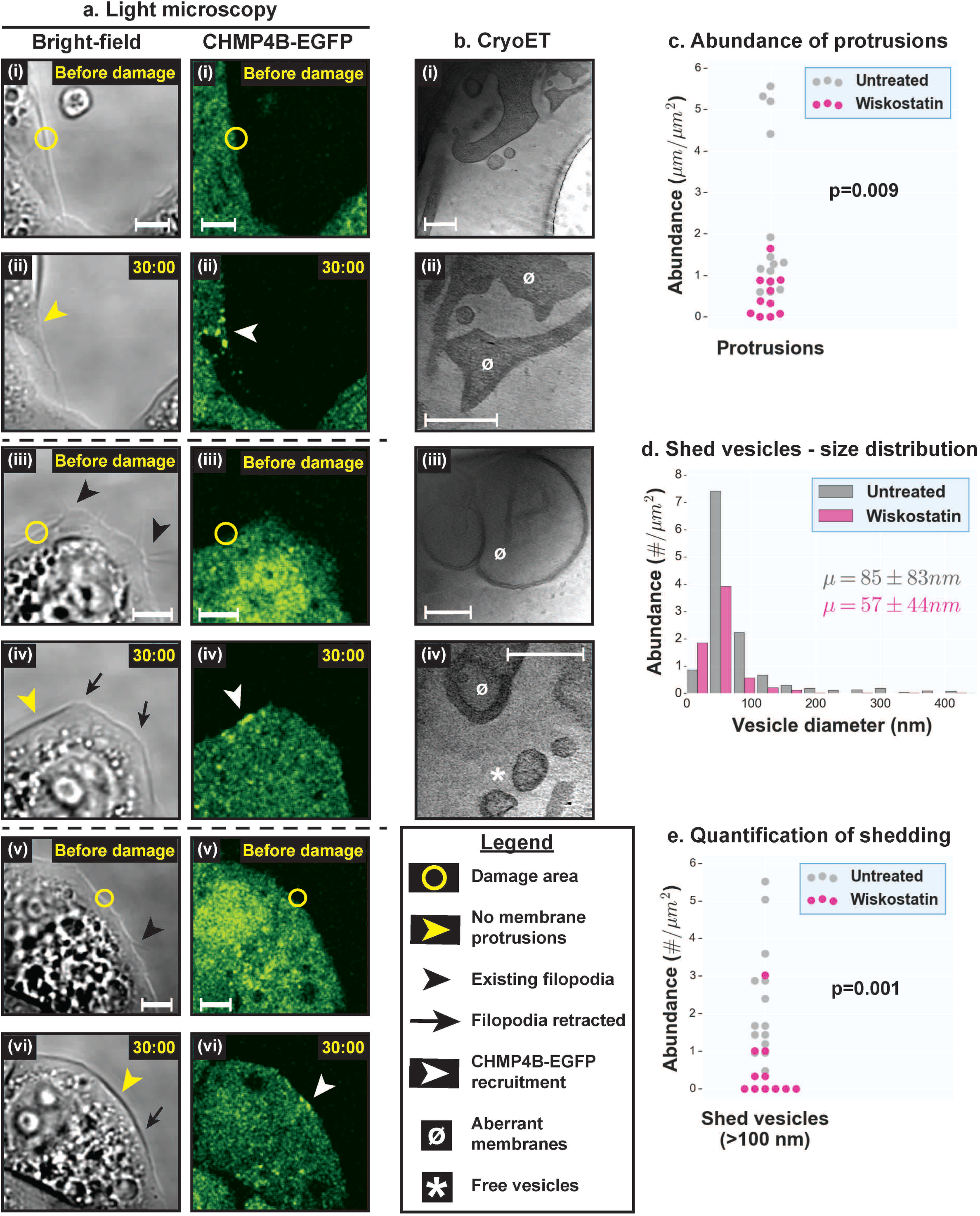
Live-cell light microscopy and cryoET of damage sites in the presence of Wiskostatin. (a) Light microscopy images of three different HeLa cells treated with wiskostatin on glass, (i, iii, v) before and (ii, iv, vi) after damage – (left panels) bright-field, and (right panels) CHMP4B-EGFP imaging. The damage areas are 3 µm in diameter. (b) CryoET of damage sites in wiskostatin-treated cells showing several aberrant membranes, a few plasma membrane protrusions and a few shed vesicles. (c) Quantification of plasma membrane protrusions at damage sites (total length of protrusions in µm per µm^2^ tomogram X-Y cross-sectional area) in wiskostatin-treated cells versus untreated cells. (d) Size distribution of shed vesicles at damage sites in wiskostatin-treated cells versus untreated cells (# vesicles in each size range per µm^2^ tomogram X-Y cross-sectional area) along with mean ± s.d. values. (e) Quantification of shed vesicles at damage sites in wiskostatin-treated cells versus untreated cells (# vesicles per µm^2^ tomogram X-Y cross-sectional area). In (c) and (e), each data point represents a tomogram and the distributions for the pairs of samples being compared are significantly different from each other as shown by KS tests. Scale bars – (a-b) 5 µm, and (c) 200 nm.

### Myo1a is involved in the organization of protrusions and/or vesicle shedding

Actin-based membrane protrusions have been previously implicated in vesicle shedding in the brush borders of gut enterocytes^26, 41^, suggesting possible similarities in the molecular machinery between the two systems. In microvilli, Myo1a forms radial densities connecting actin bundles to the plasma membrane^26^. We observed similar densities in both filopodia and damage site protrusions (Fig. 3), although they were less abundant, more irregular and harder to quantify than those described in microvilli. We therefore decided to directly test whether Myo1a plays a role in damage-mediated shedding. We knocked down Myo1a expression in cells using siRNAs and observed a significant, though not complete, reduction of Myo1a protein levels (Supplementary Fig. 4a). We then performed damage experiments on cells that showed efficient co-transfection of BLOCK-iT Alexa Fluor Red Fluorescent control RNA (to limit our analysis to transfected cells; Supplementary Fig. 4b). When these cells were laser-damaged, we observed CHMP4B-EGFP recruitment to the damage sites, with or without membrane blebbing, and loss of nearby filopodia and formation of new protrusions (Fig. 5a), just as in wildtype. By cryoET, we observed additional similarities with wildtype: (1) similar abundance of plasma membrane protrusions (Supplementary Fig. 5a); (2) similar organization of F-actin in protrusions (Fig. 5b); (3) pearling at sites of disorganized F-actin along protrusions (Fig. 5b); (4) similar abundance of shed vesicles with comparable size distribution (Supplementary Fig. 5b,c); and (5) protein densities underneath the plasma membrane in both budding profiles and free vesicles (Fig. 5b). These observations indicate that Myo1a is not absolutely essential for the organization of plasma membrane protrusions, although it is also possible that the knockdown of the protein was insufficient to see an effect or that there is significant functional redundancy with other motor proteins. We did, however, observe some defects in scission of budding vesicles (Fig. 5b – panel vi), as well as extended constrictions in protrusions (Fig. 5b – panels vii-viii) that are either disorganized membrane protrusions or defective membrane scission events. Such defects, absent in wildtype, were reflected in the wider distribution of widths for damage site protrusions (Fig. 5c; s.d. of 98 nm for Myo1a knockdown cells and 50 nm for wildtype) and a significant increase in budding profiles (and constriction events) of all sizes compared to wildtype (Fig 5d and Supplementary Fig. 5d). Therefore, Myo1a, although probably not essential, is likely involved in damage-induced shedding.

**Fig. 5:**
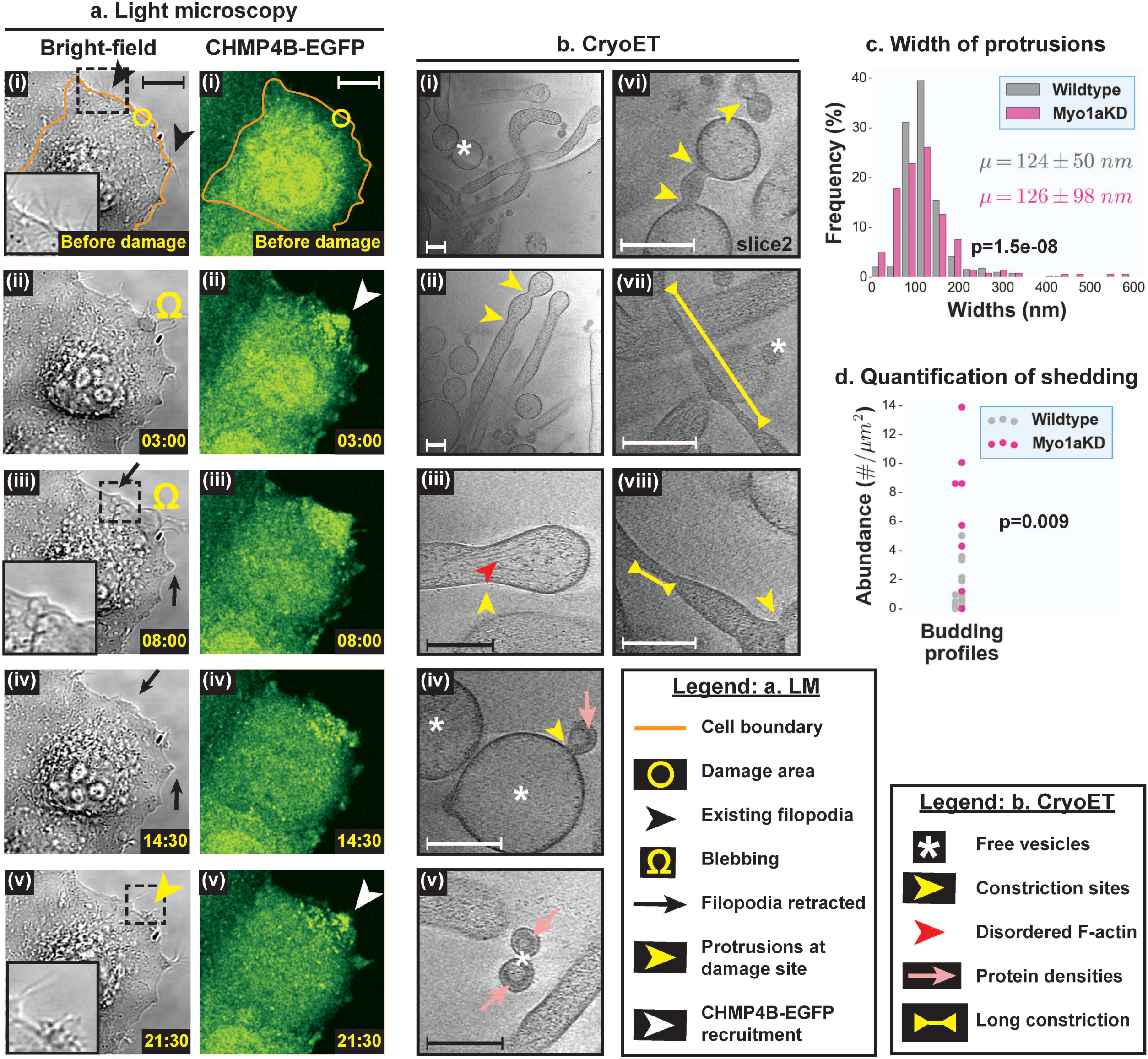
Live-cell light microscopy and cryoET of damage sites in Myo1a knockdown cells. (a) Light microscopy images of Myo1a knockdown HeLa cells grown on glass (i) before, and (ii-v) at various time points after damage – (left panels) bright-field, and (right panels) CHMP4B-EGFP imaging. The damage area is 3 µm in diameter. (b) CryoET of damage sites in Myo1a knockdown cells showing actin-filled plasma membrane protrusions, pearling/budding profiles, shed vesicles, protein densities observed at certain sites of high membrane curvature in budding profiles and shed vesicles, defective budding profiles and long constriction necks. (c) Width distribution of damage site protrusions in Myo1a knockdown cells versus wildtype (% in each size range) along with mean ± s.d. values. (d) Quantification of budding profiles at damage sites of Myo1a knockdown cells versus wildtype (# buds per µm^2^ tomogram X-Y cross-sectional area). Each data point represents a tomogram. In (c) and (d), the distributions for the pairs of samples being compared are significantly different from each other as shown by KS tests. Scale bars – (a) 10 µm, and (b) 200 nm.

### ESCRT is involved in membrane scission during shedding

ESCRT proteins are known to catalyze several membrane scission processes with similar membrane topology to shedding^42, 43^. In a previous study, recruitment of ESCRT proteins was shown to directly correlate with wound closure^2^. Furthermore, the authors observed a few extracellular membrane vesicles at sites of damage by SEM, leading to the hypothesis that ESCRT proteins close wounds by shedding damaged membranes. In this study, we observed punctate localization of CHMP4B-EGFP along membrane protrusions at damage sites. These protein foci could be localized to the necks of budding profiles, thus suggesting a role for ESCRT in membrane scission. However, it is also possible that ESCRT is instead localized to endocytic compartments such as multivesicular bodies (MVBs) and other vesicles in the protrusions. We therefore proceeded to directly test the role of ESCRT in membrane shedding.

We knocked down Vps4B, an essential AAA ATPase in the ESCRT pathway that was previously shown to be important for wound repair, and performed plasma membrane damage experiments. Knockdown of Vps4B was very efficient (Supplementary Fig. 6a) and only cells showing strong signal from co-transfected BLOCK-iT Alexa Fluor Red Fluorescent control RNA were imaged (Supplementary Fig. 6b). Again, we saw that the plasma membrane exhibited blebbing at the site of damage, existing filopodia were retracted around the damage site, new protrusions were formed at the damage site, and CHMP4B-EGFP was recruited to the damage site (Fig. 6a). CryoET of damage sites showed (1) numerous membrane protrusions, (2) F-actin bundles in protrusions, and (3) pearling (Fig. 6b and Supplementary Fig. 7a), all similar to wildtype. Although the abundance of these damage site protrusions seemed to show a different distribution compared to wildtype (as indicated by the p-value from a Kolmogorov-Smirnov test), the mean was quite similar to that of wildtype cells (∼2.5 µm of protrusions per µm^2^ of tomogram X-Y cross-sectional area for both samples; Supplementary Fig. 7a). These observations indicate that Vps4B is not essential for the organization of membrane protrusions. However, there was an appreciable decrease in the number of shed vesicles smaller than 200 nm in diameter compared to wildtype (Supplementary Fig 7b), as indicated by the mean vesicle diameter (141 ± 141 nm for Vps4B knockdown cells compared to 85 ± 83 nm for wildtype; mean ± s.d.). There was a corresponding increase in the abundance of budding profiles (or constriction events) smaller than 200 nm in diameter (Supplementary Fig. 7c). These smaller budding profiles sometimes formed long chains (Fig. 6b – panel v) reminiscent of failed HIV-1 budding profiles from the plasma membrane upon disruption of Vps4 function^44^. Both budding profiles and smaller shed vesicles displayed protein densities underneath their membrane, as seen in wildtype, and the densities were enriched in the chain of budding profiles (Fig. 6b – panel v). In addition to wildtype-like protrusions, we occasionally observed shed protrusions devoid of F-actin (Fig. 6b – panel vi) and a few nested protrusions with or without F-actin (Fig. 6b – panels vii-ix). These defects, not seen in wildtype, are likely due to accumulation of membrane from unshed vesicles upon disruption of Vps4 function and are quantitatively reflected in the greater spread of protrusion widths (Supplementary Fig. 7d; s.d. of 98 nm for Vps4B knockdown cells compared to 50 nm for wildtype). Vps4B is therefore likely involved in membrane scission in damage-induced shedding, particularly for vesicles smaller than 200 nm in diameter, consistent with its published role in closure of small wounds^2^.

**Fig. 6:**
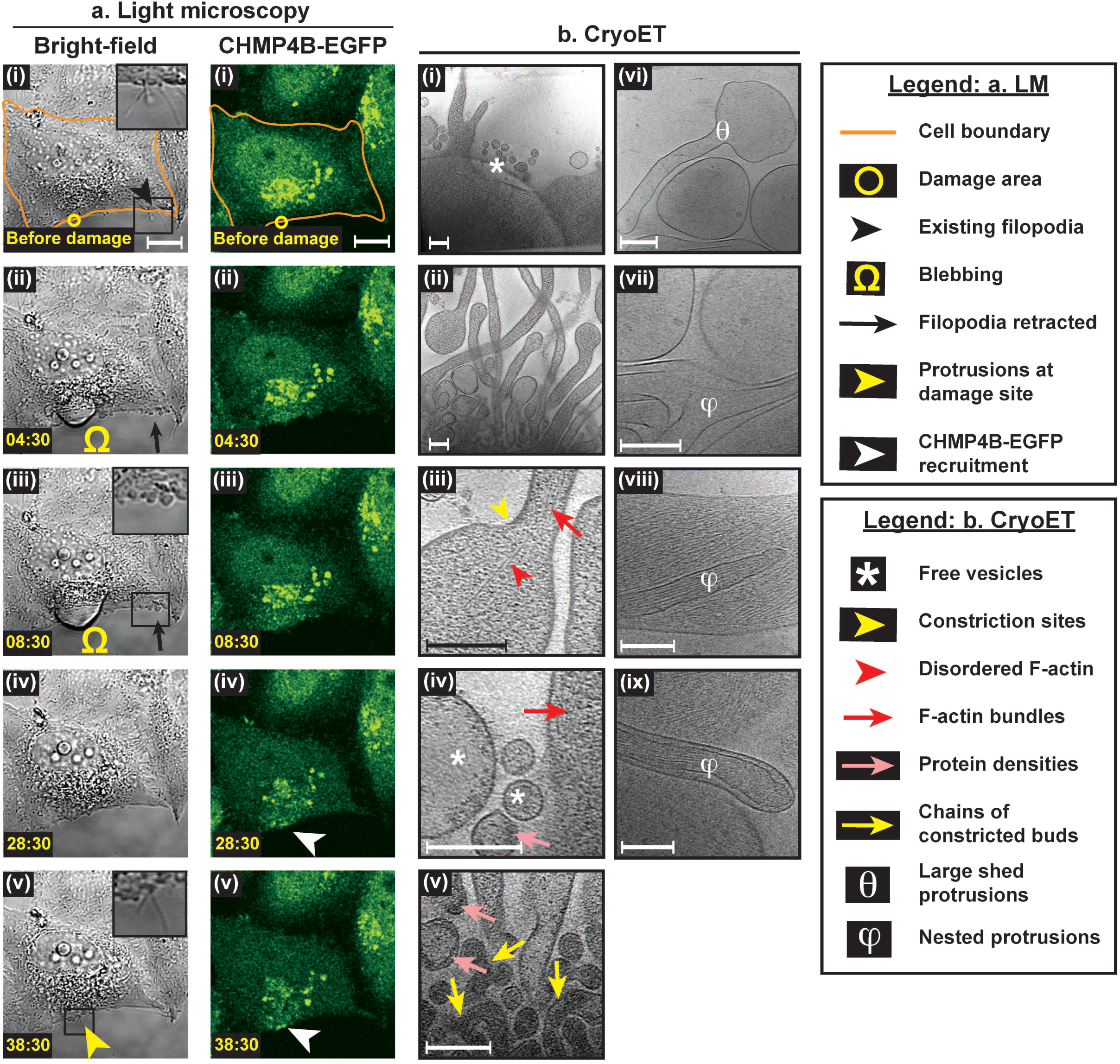
Live-cell light microscopy and cryoET of damage sites in Vps4B knockdown cells. (a) Light microscopy images of Vps4B knockdown HeLa cells grown on glass (i) before, and (ii-v) at various time points after damage – (left panels) bright-field, and (right panels) CHMP4B-EGFP imaging. The damage area is 3 µm in diameter. (b) – CryoET of damage sites of Vps4B knockdown cells showing actin-filled plasma membrane protrusions, pearling/budding profiles, shed vesicles, protein densities observed at certain sites of high membrane curvature in budding profiles and shed vesicles, chains of budding profiles, shed membrane protrusions devoid of F-actin and nested protrusions. Scale bars – (a) 10 µm, and (b) 200 nm.

## Discussion

Here we used correlated light microscopy and cryoET to explore plasma membrane damage repair in HeLa cells. Our findings lead to a model summarized in Fig. 7: (1) actin and membrane from neighboring regions of the cell are relocated to the site of damage; (2) in a process dependent on F-actin nucleation and Myo1a, relocated membrane and actin are used to construct new filopodia-like protrusions to act as scaffolds for vesicle shedding; and (3) F-actin dynamics, Myo1a and the ESCRT machinery mediate membrane remodeling and scission to shed damaged membrane. Damage-induced plasma membrane shedding is thus more complex than current models depicting simple vesiculation from flat plasma membrane domains^13, 45^. Interestingly, tufts of microvilli-like plasma membrane protrusions were previously reported in bovine retinal microvascular endothelial (BRME) cells in response to wounding^5^, and filopodia-like protrusions were observed in epithelial cells of *Drosophila* embryos upon wounding and were demonstrated to be important for healing^46^. However, the possible function of these structures in membrane shedding remained unknown until now.

**Fig. 7:**
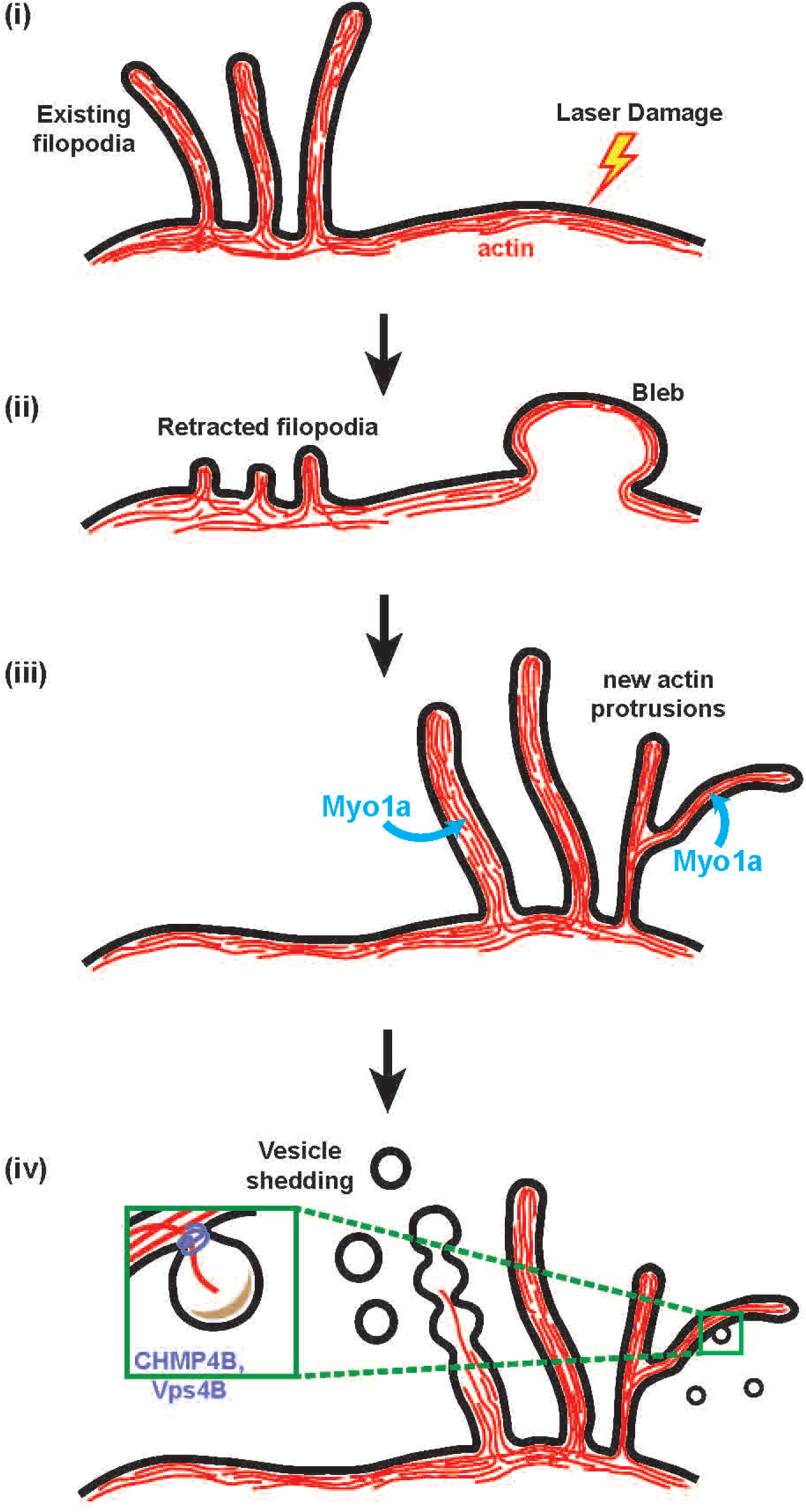
Model for damage-induced plasma membrane shedding. (i-ii) Actin and membrane are redirected to the site of damage from other regions of the cell (including neighboring filopodia), often resulting in the formation of membrane blebs. (iii) Membrane blebs are withdrawn into the cell and reorganized into membrane protrusions using F-actin. (iv) F-actin depolymerization could cause membrane deformation and pearling. Proteins that sense or induce high negative membrane curvature could provide additional force necessary for membrane deformation (especially for smaller vesicles). Myo1a is involved in building protrusions and/or vesicle shedding while Vps4B is involved in membrane scission and shedding, especially of smaller vesicles.

In this study, we observed shedding from membrane protrusions even 10-15 min post-damage, although wound closure has been reported to occur within a couple minutes of damage. It is therefore possible that wound closure involves other mechanisms before shedding discards membrane patches to complete the repair process. For example, reassembly of the actin cortex or recruitment of annexins to the damage site could maintain a diffusion barrier until these membrane patches are shed. Consistent with this hypothesis, annexin A1 and annexin A6 have been previously reported to be enriched in plasma membrane protrusions and shed vesicles upon permeabilization of HEK 293 cells with the toxin SLO^11^.

In the first step of shedding, membrane could be transferred to sites of damage by lateral diffusion or by a more complex process involving endocytosis at the source followed by exocytosis at the sites of damage. We observed numerous internal vesicles near damage sites in the cytosol and inside protrusions, supporting the endocytosis-exocytosis model. Retraction of existing filopodia at nearby sites suggests that they contribute actin and membrane for remodeling of damage sites. To our knowledge, a function for filopodia as a reservoir for membrane and actin during plasma membrane repair has not been suggested previously.

Damage site protrusions may well be repurposed filopodia. They share several features (summarized in results), and are distinguished mainly by the increased shedding from the former. Strikingly, we found that they even share the molecular marker Pls1 (an actin-bundling protein). Consistent with this model, even canonical filopodia exhibit basal levels of shedding, although this function has not been well studied. Filopodia-like protrusions could offer an advantage over a flat patch of plasma membrane by providing a higher membrane curvature more suitable for shedding. As already noted, shedding from brush-border enterocytes occurs from microvilli, another form of plasma membrane protrusions^26, 41^. Another study reported shedding from mesenchymal cells in response to progesterone treatment via putative vesicle-budding events on plasma membrane protrusions^47^. Thus, shedding from actin-rich membrane protrusions could be a universal mechanism.

Damage site filopodia are likely built by F-actin along with Myo1a and other motor proteins. As reported for canonical filopodia^38^, we found several free ends of F-actin throughout the length of all filopodia (including ones at damage sites), supporting a de novo filament nucleation model that describes filopodia growth through multiple nucleation events along its length. Growing F-actin at the tip could provide the force to propel membrane forward while the polymerized F-actin bundles dictate the shape of the filopodia; for example, in curved filopodia we always saw F-actin filaments/bundles of a similar contour closely associated with their plasma membrane (Fig. 3a – panel v). It remains to be seen how different actin nucleation pathways (Arp2/3 vs. Formins) contribute to the formation of damage site filopodia since they show both branched F-actin at their base (consistent with the role of Arp2/3) and long linear F-actin filaments along their length (consistent with Formins). F-actin growth in filopodia could be accompanied by Myo1a or other motor proteins (including Myo1b and Myo6) moving membranes on these filament-tracks towards the tip; Myo1b was reported to localize to filopodia^48^ and an engineered form of Myo6 was shown to induce formation of filopodia^49^, both in HeLa cells. All three motor proteins have been reported to bind phosphoinositides (PIs), which are enriched in filopodia. In addition to plasma membrane moved along F-actin by myosin motors, cytosolic vesicles trafficked internally along F-actin within the filopodia could be used to meet the membrane demands of a growing filopodium. Such internal vesicles, which we observed in damage site filopodia, could also help sort damaged membrane domains to the sites of shedding.

The final steps of shedding involve membrane deformation and scission in order to release free vesicles to the exterior. It is worth noting that the free vesicles observed in this study showed a very broad size distribution. This size variation and the presence of budding profiles on damage site filopodia argue against the possibility that these vesicles are exosomes (derived from fusion of MVBs with the plasma membrane); exosomes are smaller (<80 nm in diameter) and more or less uniform in size^13^.

We postulate two different mechanisms, not mutually exclusive, that could cause deformation of membrane domains for shedding: (1) rapid depolymerization of F-actin could cause unsupported membrane to vesiculate or pearl; and (2) membrane-binding proteins could sense/induce high negative curvature, particularly for smaller vesicles. Several previous reports support our first model: (1) destruction of the F-actin cortex with latrunculin A induces pearling of plasma membrane^50^; (2) actin polymerization, destabilization of the actin cytoskeleton and disruption of actin-membrane interactions induce shedding from neutrophils and platelets^51–53^; and (3) microvilli of enterocytes that shed vesicles from their tip show a similar lack of tubular morphology at their tips that correlates with a lack of F-actin bundles in these regions^26, 41^. In support of our second model, we observed protein densities underneath membrane regions of high curvature in budding profiles and smaller vesicles. Several inverse BAR (I-BAR) proteins such as IRSp53 have been reported to have high affinity for PIs (enriched in filopodia) and induce negative curvature during various kinds of cellular morphogenesis including filopodia formation^54, 55^. Moreover, they are known to couple membrane deformation to actin dynamics, an important feature of shedding in our experiments. Involvement of these BAR-domain proteins would not come as a surprise since one such protein, Angiomotin, is known to function in another process involving negative membrane curvature, namely HIV-1 budding from the host cell^56^.

Subsequent to membrane deformation, the ESCRT machinery is involved in membrane scission during shedding. Previous work hypothesized this function for ESCRT due to their involvement in wound closure and several other membrane scission processes of similar topology^2^. Here we show direct evidence for this function by showing a defect in membrane scission (accumulated budding profiles and a reduced number of free vesicles <200 nm in diameter) when Vps4B function is disrupted. This defect was not complete, so there is likely functional redundancy in the ESCRT system, perhaps with Vps4A.

## Methods

### Cell growth

A HeLa Kyoto cell line stably expressing CHMP4B-EGFP (gift from Dr. Anthony A. Hyman, Max Planck Institute) was grown in a humidified 37 °C incubator with a constant supply of 5% CO_2_. Cells were cultured in high glucose (L-Glutamine +) Dulbecco’s modified Eagle’s medium (DMEM; DML09, Caisson Labs, Smithfield, UT) supplemented with 10% fetal bovine serum (FBS; Cat No – 10437028, ThermoFisher), 1 mM sodium pyruvate (Cat No – 11360070, ThermoFisher), 100 units/mL penicillin and 100 µg/mL streptomycin. The CHMP4B-EGFP plasmid was maintained in these cells using 400 µg/mL G-418 disulfate (Cat No – G64500, Research Products International). For experiments involving confocal microscopy, cells were grown on poly-D-lysine-coated 35 mm coverslip bottom dishes (P35GC-1.5-14-C, MatTek corporation). For experiments further involving CLEM and cryoET, cells were grown on 200 mesh gold R2/2 London Finder Quantifoil grids (Quantifoil Micro Tools GmbH, Jena, Germany) added to the bottom of MatTek dishes. Prior to addition of these grids to the MatTek dishes, they were coated with 0.1 mg/mL human fibronectin (Cat No – C-43060, PromoCell) by floating them on fibronectin droplets on parafilm for approx. 15-30 min. Additionally, they were coated with 10 nm Au fiducials to be later used for tomography. Roughly 4 µL of 15 X diluted Au fiducials (Cat No – 15703, Ted Pella) in 0.01% bovine serum albumin (BSA) were dried onto the grids. Cells were grown to a density of approx. two to three per grid-square over a period of one to three days depending on the experiment.

### Gene silencing, expression of fluorescent proteins and drug treatments

Knockdown experiments for Myo1a and Vps4B were performed in CHMP4B-EGFP expressing HeLa cells using Lipofectamine RNAiMax (Cat No – 13778075, ThermoFisher). Cells were grown on 35mm MatTek dishes or on grids placed at the bottom of these dishes overnight. They were transfected with 50 pmol of siMyo1a (SMARTpool ON-TARGETplus MYO1A siRNA, Cat No – L-008765-01, GE Dharmacon) or with 50 pmol of siVps4B (ON-TARGETplus Human VPS4B siRNA, Cat No – L-013119-00, GE Dharmacon). Transfections were performed for two rounds, each lasting for 24 hours. Cells were co-transfected with BLOCK-iT Alexa Fluor Red Fluorescent Control (Cat No – 14750100, ThermoFisher) to ensure transfection efficiency and to identify transfected cells for photo-damage experiments. Transfection experiments with RFP-LifeAct and FusionRed-Fimbrin/Plastin-1 (Pls1) were conducted similar to other transfections before live cell microscopy. For experiments involving Wiskostatin (Cat No – 4434, Tocris Bioscience), the drug was administered to the cells at 25 µM for 2-3 hours prior to photo-damage experiments.

### Confocal microscopy and laser damage

Imaging was performed at the Caltech Biological Imaging Facility on a Zeiss LSM800 microscope equipped with a large environmental chamber to maintain the temperature at 37 °C and a smaller insert module that helped maintain both the temperature and a CO_2_ level of 5%. Prior to confocal microscopy and laser damage, the photosensitizer Al(III) Phthalocyanine chloride disulfonic acid (AlPcS2a; Cat No – P40632, Frontier Scientific) was added to the cell medium at ∼1.3 µM final concentration. Laser damage experiments were performed within the next 10 min to prevent any large interference from endocytosed photosensitizer. No washes were performed after incubation with the photosensitizer. The cell media also contained 50 mM HEPES (Cat No – 15630080, ThermoFisher) to prevent pH fluctuations during cell transport and handling. Both bright-field and fluorescence imaging were performed using an LD C-Apochromat 40 X water-immersion objective with an NA of 1.1 and images were recorded using Photo Multiplier Tubes (PMTs; for bright-field image) and GaAsP-PMT (for fluorescence). Green fluorescence imaging was performed using a diode laser at 488 nm at ∼1.5-2% of its maximum power. Photo-damage was administered using a diode laser at 640 nm operated at 100% of its maximum power. The maximum power for the laser lines was 500 mW at the source but measured to be ∼750 µW at the level of the objective lenses for the 488 nm laser and ∼400 µW for the 640 nm laser. We observed that cells grown on EM grids were more susceptible to laser damage than ones grown on glass. Therefore, we reduced the number of laser pulse cycles and the damaged area for cells on EM grids accordingly. For photo-damaging cells grown on glass, a circular area of 3 µm diameter was chosen close to the cell periphery and scanned for 100 or 150 cycles. For cells on grids, a circular area of 1.5 µm diameter was scanned for 35 cycles. Damage response and recovery were monitored intermittently (∼ every 1-2 min) for up to 1 hour after photo-damage. A scan speed of 7 was used for both photo-damaging and imaging cells. The pixel size for imaging was set at 0.312 µm (0.156 µm at a zoom factor of 2) while the image sizes were fixed at 512 × 512 pixels. For CLEM and cryoET experiments, cells were fixed at 10-15 min post-damage for 45 min with 4% paraformaldehyde (PFA; Cat No – RT-15710, Electron Microscopy Sciences) in PBS. Cells were washed 3 times with PBS before plunge-freezing for CLEM and cryoET. While studying damage response in cells transfected with siRNAs, candidate cells were chosen based on cytosolic levels of a co-transfected fluorescent BLOCK-iT RNA.

### Plunge-freezing

EM grids containing photo-damaged and fixed cells were plunge frozen in a liquid ethane/propane mixture using a Vitrobot Mark IV (FEI, Hillsboro, OR)^57^. The Vitrobot was set to 95-100% relative humidity at 37 °C and blotting was done manually from the back side of the grids using Whatman filter paper strips. Plunge-frozen grids were subsequently loaded into Polara EM cartridges (FEI) or Krios autogrid cartidges (ThermoFisher). EM cartridges containing frozen grids were stored in liquid nitrogen and maintained at ≤−170 °C throughout storage, transfer and cryo-EM imaging.

### Correlative light and electron microscopy (CLEM) and electron cryotomography (cryoET)

Cells previously photo-damaged and imaged by confocal microscopy were imaged by cryo-EM using either an FEI G2 Polara 300 kV FEG cryo-TEM or a ThermoFisher Krios G3i 300 kV FEG cryo-TEM at the Caltech CryoEM Facility. Both these microscopes were equipped with a 4k × 4k K2 Summit direct detector (Gatan, Inc.) operated in electron counting mode. An energy filter was used to increase the contrast at both medium and higher magnifications with a slit width of 50 eV and 20 eV, respectively. Additionally, defocus values of close to negative 100 and negative 8 µm were used to boost the contrast (in the lower spatial resolution range) at the medium and higher magnifications respectively. Magnifications typically used on the Polara were 3,000 X and 22,500 X (in the medium and higher ranges), corresponding to pixel sizes of 3.7 nm and 4.8 Å respectively. On the Krios, 3600 X/4800 X and 26000 X were used in the two magnification ranges that correspond to pixel sizes of 4.2 nm/3.1 nm and 5.38 Å respectively. A Volta phase plate (VPP) was optionally used on the Krios to further improve contrast at higher magnifications in certain cases. SerialEM software^58^ was used for all imaging.

Photodamaged cells were located in the electron microscope using the markers on the Finder grids. The markers were clearly visible by transmission light microscopy but only partially identifiable by cryo-EM after freezing. However, a full grid montage at a low magnification of close to 100 X is sufficient to positively identify these markers based on their overall arrangement on the grid. The photo-damaged locations in the cells were located by roughly correlating the light microscopy images with the EM maps based on the shape of the cells or by using the image registration protocol in SerialEM. Cracks, regularly spaced 2 µm holes in the carbon film, ice contamination and other features visible by both light and electron microscopy were sufficient to obtain an accurate enough correlation (< 500 nm precision) for the purpose of tomography. Once the areas of interest were identified and marked, anchor maps were used to revisit these locations and collect tilt-series in an automated fashion. Each tilt-series was collected from negative 60° to positive 60° with an increment of 1° or 2° in an automated fashion using the low dose functions of tracking and focusing. The cumulative dose of each tilt-series ranged between 80 and 150 e^-^/Å^2^. Once acquired, tilt-series were binned into 1k × 1k arrays before alignment and reconstruction into 3D tomograms with the IMOD software package^59^ and tomo3D^60^. Tilt series were aligned using 10 nm Au fiducials or patch tracking in IMOD while reconstructions were performed using SIRT in tomo3D. In addition to tilt-series, projection images were saved at other magnifications like 360 X for correlation post-data acquisition.

For data processing, analysis and generating figures, more precise correlations were performed using custom Python scripts in a semi-automated fashion. Features used as control points here were similar to the ones used for correlation during data acquisition. However, the number of control points used was larger and a robust best-fit method was employed to increase the precision for correlation. Control points that gave the most accurate correlation (based on the overall error in the fit) were selected from the set provided by the user for accurate correlation. Precision for correlation at lower magnifications is particularly important because of the large pixel sizes involved.

### Segmentation

Segmentations of tomograms were manually performed using Amira (ThermoFisher). Animations of segmented tomograms were created using Amira and Adobe Photoshop CC (Adobe Inc., San Jose, USA). Segmentations were done to the best of our abilities bearing in mind the limitation of the missing wedge of information in cryoET. Distinctions between plasma membrane and shed vesicles were based only on unambiguously segmented data.

### Quantification

#### Live-cell light microscopy

Cells were observed for ∼45 minutes post damage (or without damage). Damage sites showing any newly visible filopodia were counted positive for appearance of new protrusions (these protrusions were much more abundant at damage sites than control sites). Integrated intensities over a square area of pixels were plotted over time for damage sites and two other sites from the same cell as control. Intensity data was extracted from live-cell microscopy movies over these square areas using Fiji (ImageJ) before plotting them in Python 3.5 using Numpy and Matplotlib.

#### Actin

F-actin was analyzed post-segmentation in Amira using the filament module. Actin filaments that were unambiguously linear (no branching, showing free ends on either side) were used for length measurements. A total of ∼450 filaments were measured for filopodia and ∼480 filaments for damage site protrusions. Inter-filament spacing was measured manually at multiple positions between parallel F-actin filaments using IMOD. A total of ∼150 measurements were made for filopodia and ∼100 for damage site protrusions.

#### Other measurements from cryoET

Quantifications were performed on a per-tomogram basis. For each damage site, 1-2 tomograms were randomly selected close to the site of CHMP4B-EGFP recruitment. The following numbers of tomograms were selected for each sample – 14 tomograms from wildtype damage sites, 10 from control sites with no damage, 11 from damage sites of wiskostatin-treated cells, 8 from damage sites of Myo1a knockdown cells and 11 from damage sites of Vps4B knockdown cells. All measurements were made using IMOD. Density of protrusions was measured as total lengths of protrusions in a tomogram (in µm) divided by tomogram X-Y cross-sectional area (in µm^2^). Widths of plasma membrane protrusions were measured at regular intervals at local maxima, minima and anywhere in between. Density of shed vesicles was measured as number of vesicles per µm^2^ cross-sectional per tomogram. The vesicle sizes were measured as cross-sectional diameters. Similar quantifications were done for budding profiles as well. New budding profiles in shed vesicles were included in the analysis. Internal vesicles were measured as number of vesicles per µm length of protrusion per tomogram. All model files were exported as text before plotting using Numpy and Matplotlib libraries in Python 3.5. Kolmogorov-Smirnov tests (KS tests) were performed using the ttest_ind method from the Scipy package. We chose one-tailed t-tests for our statistics because our EM data clearly indicated a one-sided shift in the parameter being measured.

## Acknowledgements

This work was supported by funding from the NIH (P50 AI150464 awarded to G.J.J.). We thank Dr. Anthony A. Hyman and Dr. Ina Poser for providing the HeLa cell line stably expressing CHMP4B-EGFP. We thank Dr. Andres Collazo and Steven Wilbert for technical assistance with confocal microscopy. We also thank Dr. Songye Chen and Dr. Andrey Malyutin for technical assistance with electron cryomicroscopy. The bulk of the confocal imaging was performed at the Biological Imaging Facility and electron microscopy was performed at the Beckman Institute Resource Center for Transmission Electron Microscopy, both at Caltech.

## Supplementary Material

**Supplementary Fig. 1: Live-cell light microscopy of plasma membrane damage site showing pearling.**

Light microscopy images of HeLa cells grown on glass (i) before, and (ii-iv) at various time points after damage showing membrane pearling – (left panels) bright-field images, (middle panels) magnified images of insets in the left panels, and (right panels) CHMP4B-EGFP images. The damage area is 3 µm in diameter. Scale bars – 10 µm.

**Supplementary Fig. 2: Live-cell light microscopy of plasma membrane damage site showing retraction fibers.**

Light microscopy images of HeLa cells grown on glass (i) before, and (ii-iii) at various time points after damage – (left panels) bright-field images showing retraction fibers, and (right panels) CHMP4B-EGFP images showing its recruitment to the retraction fibers. The damage area is 3 µm in diameter. Scale bars – 10 µm.

**Supplementary Fig. 3: Schematic for the experimental CLEM workflow.**

**Supplementary Fig. 4: Efficiency of siRNA transfection and Myo1a knockdown.**

(a) (i) Western blot showing the protein levels of Myo1a upon siRNA-mediated knockdown compared to control siRNA transfected cells. Actin serves as the loading control. (ii) Quantification of Myo1a protein levels from three different knockdown experiments relative to control siRNA transfection experiments and normalized using actin loading control. s.d. values are shown as error bars. (b) Light microscopy images of damaged cell in Fig. 5a – (left) bright-field image, (middle) CHMP4B-EGFP expression, and (right) BLOCK-iT-AlexaFluor-Red control RNA; the latter used as a readout for transfection efficiency.

**Supplementary Fig. 5: Damage response in Myo1a knockdown cells by cryoET.**

(a) Quantification of plasma membrane protrusions at damage sites of Myo1a knockdown cells versus wildtype (total length of protrusions in µm / µm^2^ tomogram X-Y cross-sectional area). (b) Quantification of shed vesicles at damage sites of Myo1a knockdown cells versus wildtype (# vesicles / µm^2^ tomogram X-Y cross-sectional area). (c) Size distribution of shed vesicles at damage sites of Myo1a knockdown cells versus wildtype (# vesicles in each size range / µm^2^ tomogram X-Y cross-sectional area) along with their mean ± s.d. values. (d) Size distribution of budding profiles at damage sites of Myo1a knockdown cells versus wildtype (# budding profiles in each size range / µm^2^ tomogram X-Y cross-sectional area) along with their mean ± s.d. values. In (a) and (b), each data point represents a tomogram and the distributions for the pairs of samples being compared are not significantly different from each other as shown by KS tests.

**Supplementary Fig. 6: Efficiency of siRNA transfection and Vps4B knockdown.**

(a) (i) Western blot showing a reduction in the protein levels of Vps4B upon siRNA-mediated knockdown compared to control siRNA transfected cells. Actin serves as the loading control. (ii) Quantification of Vps4B protein levels from three different knockdown experiments relative to control siRNA transfection experiments and normalized using actin loading control. s.d. values are shown as error bars. (b) Light microscopy images of damaged cell in Fig. 6a – (left) bright-field image, (middle) CHMP4B-EGFP expression, and (right) BLOCK-iT-AlexaFluor-Red control RNA; the latter used as a readout for transfection efficiency.

**Supplementary Fig. 7: Damage response in Vps4B knockdown cells by cryoET.**

(a) Quantification of plasma membrane protrusions at damage sites of Vps4B knockdown cells versus wildtype (total length of protrusions in µm / µm^2^ tomogram X-Y cross-sectional area). Each data point represents a tomogram. The two distributions are significantly different from each other as shown by KS test. However, their means are very similar (∼2.5 µm of protrusions per µm^2^ tomogram X-Y cross-sectional area). (b) Size distribution of shed vesicles at damage sites of Vps4B knockdown cells versus wildtype (# vesicles in each size range / µm^2^ tomogram X-Y cross-sectional area) along with their mean ± s.d. values. (c) Size distribution of budding profiles at damage sites of Vps4B knockdown cells versus wildtype (# buds in each size range / µm^2^ tomogram X-Y cross-sectional area) along with their mean ± s.d. values. (d) Width distribution of protrusions at damage sites of Vps4B knockdown cells versus wildtype (% in each size range) along with their mean ± s.d. values. In each of the pairwise comparisons, the distributions for the two samples were significantly different from each other as indicated by the p-values of KS tests.

**Supplementary Movie 1: Live-cell light microscopy of laser-damaged cells (6 cells, including the one shown in Fig. 1).**

**Supplementary Movie 2: Representative experiment including CLEM, cryoET and segmentation of a damage site.**

**Supplementary Movie 3: Live-cell light microscopy of a laser-damaged cell showing FusionRed-Pls1 localization in both regular filopodia and damage-induced protrusions.**

The damage area (yellow circle) is 3 µm in diameter and the time points after damage are denoted in minutes:seconds.

