## Supplementary Figures for "Plasma membrane damage removal by F-actin-mediated shedding from repurposed filopodia"

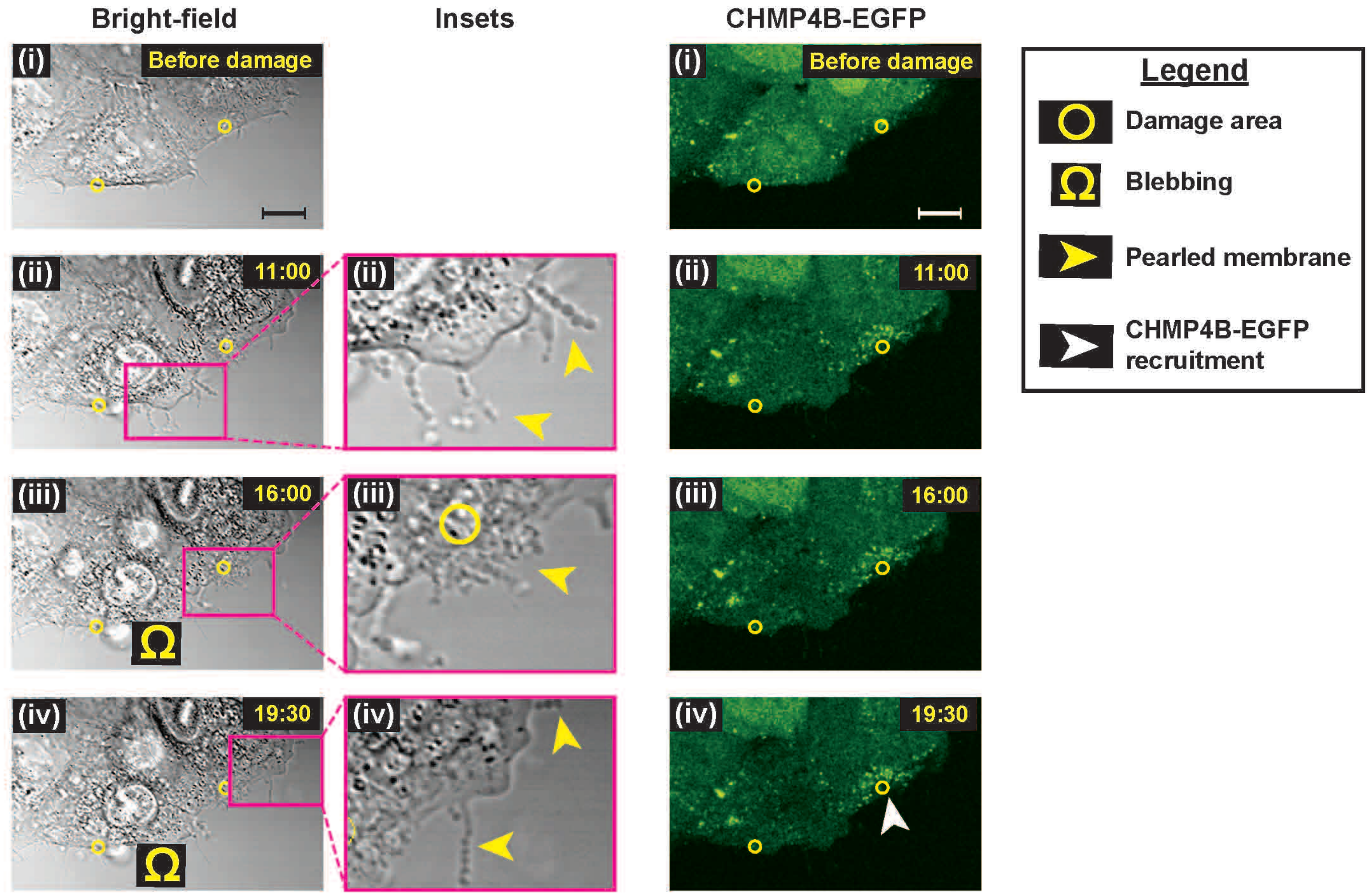


**Supplementary Fig. 1: Live-cell light microscopy of plasma membrane damage site showing pearling.**

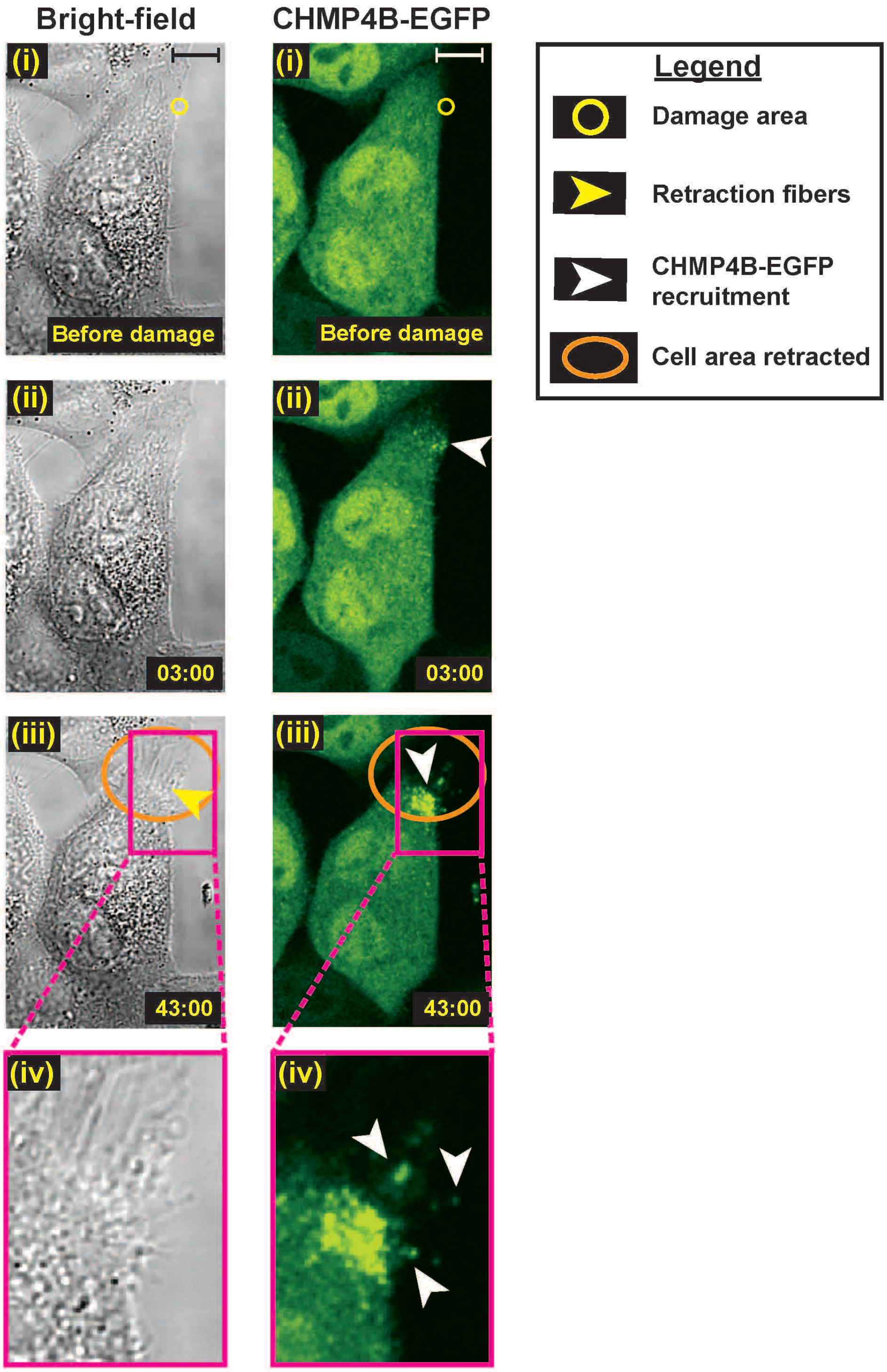


**Supplementary Fig. 2: Live-cell light microscopy of plasma membrane damage showing retraction fibers.**

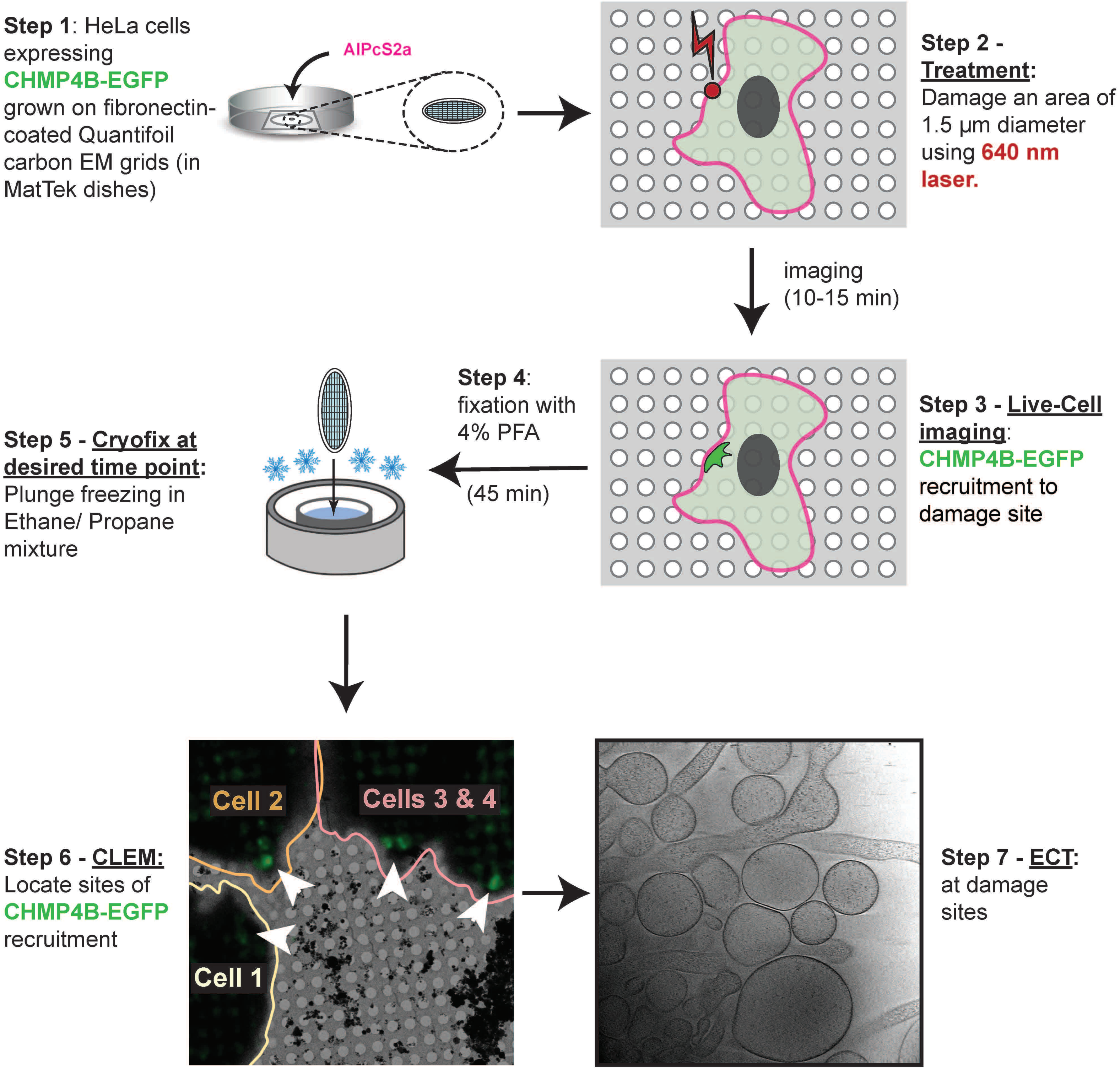


**Supplementary Fig. 3: Schematic for the experimental workflow.**

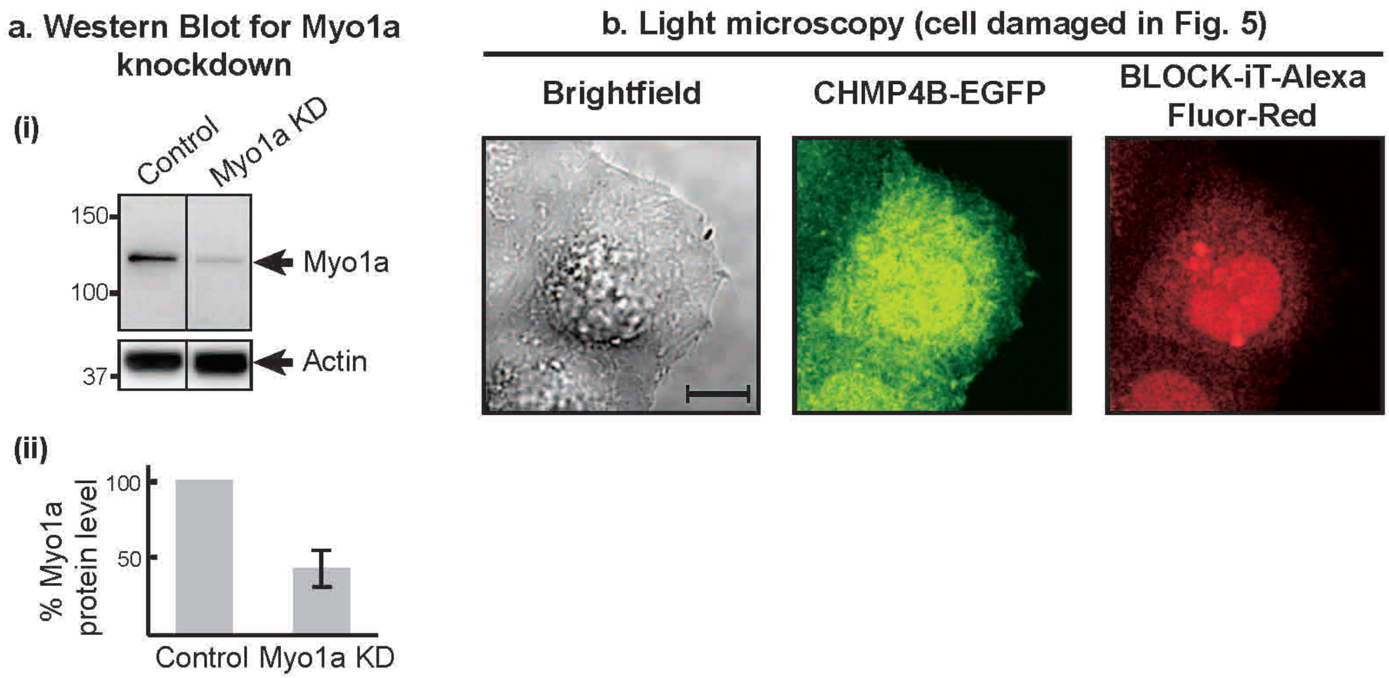


**Supplementary Fig. 4: Efficiency of siRNA transfection and Myo1a knockdown.**

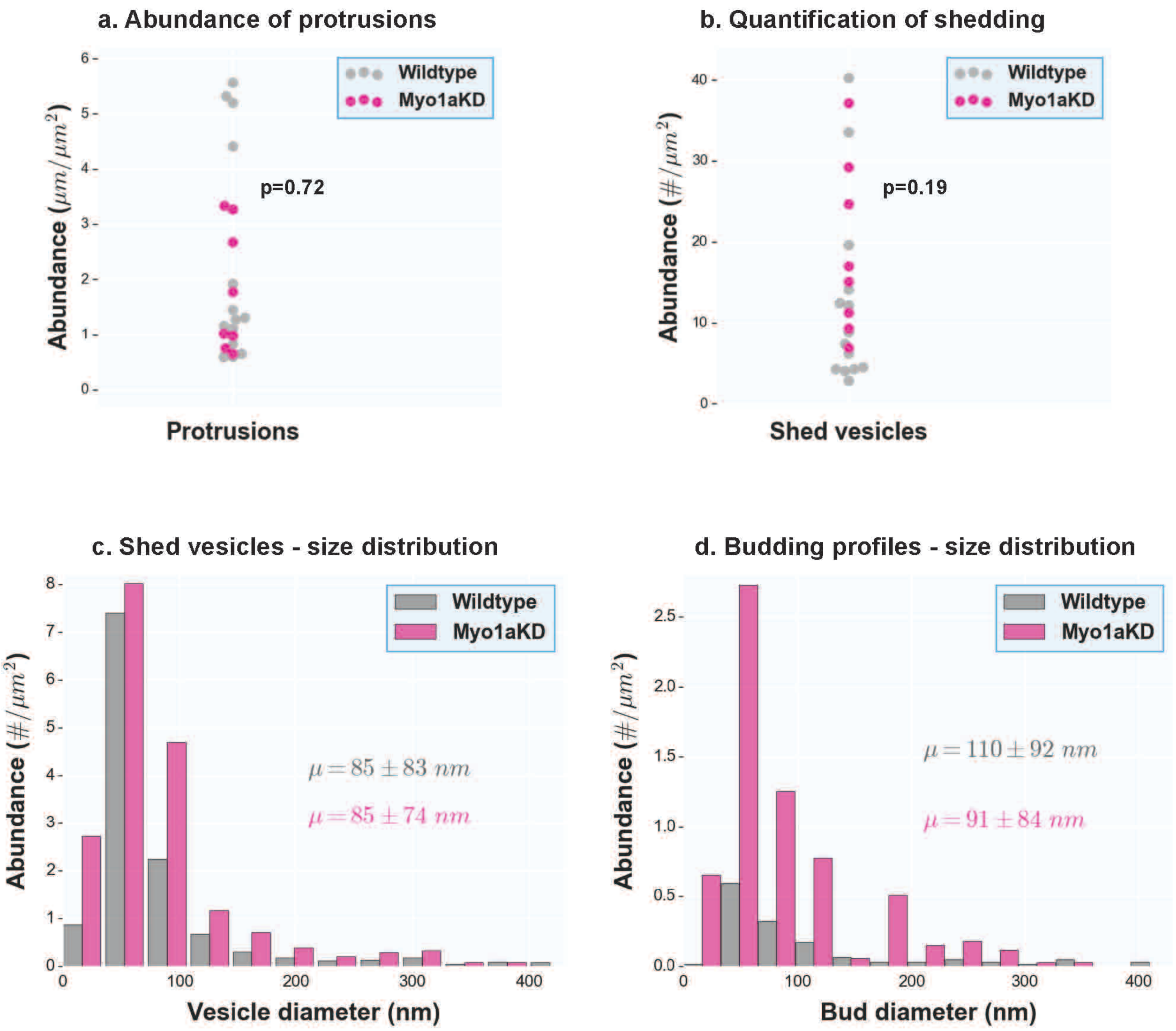


**Supplementary Fig. 5: Damage response in Myo1a knockdown cells by cryoET.**

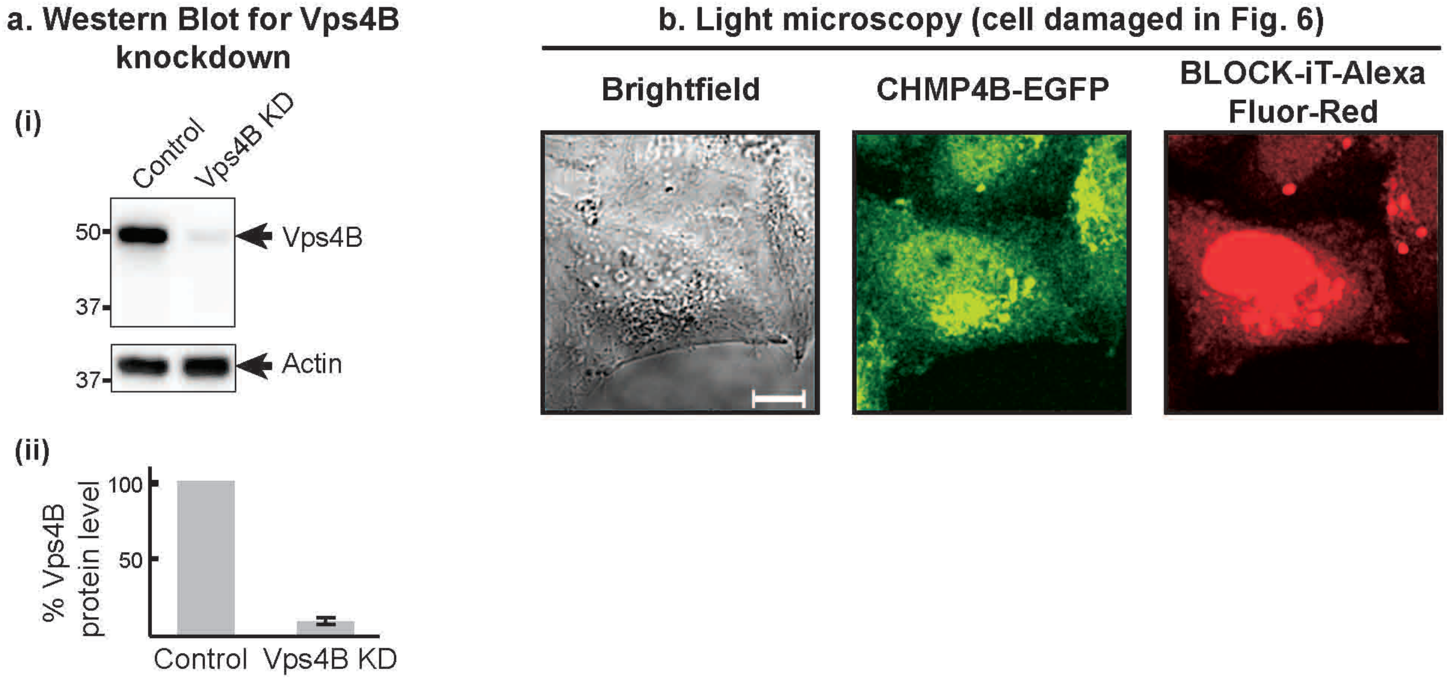


**Supplementary Fig. 6: Efficiency of siRNA transfection and Vps4B knockdown.**

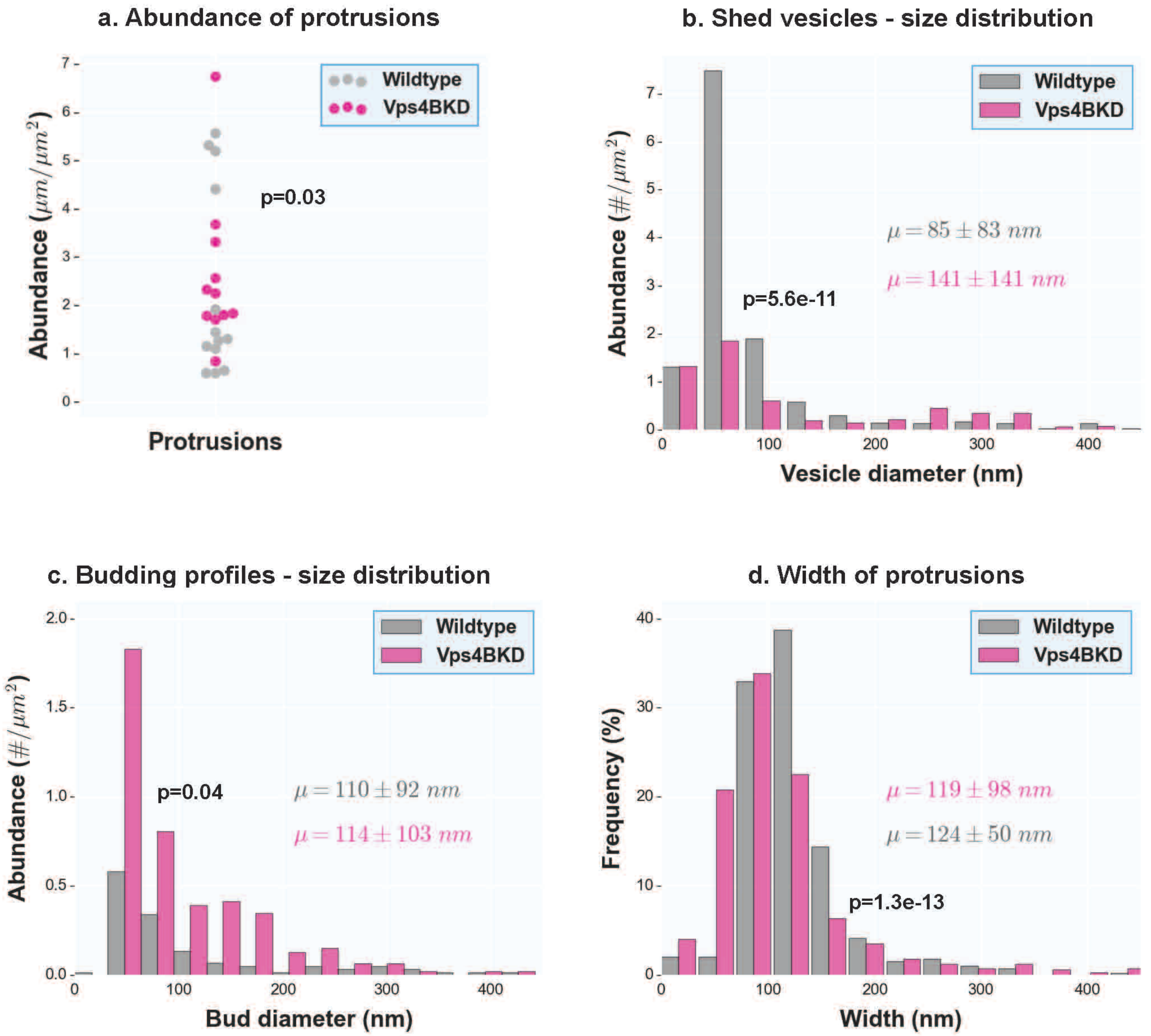


**Supplementary Fig. 7: Damage response in Vps4B knockdown cells by cryoET.**
